# Synthesis of phylogeny and taxonomy into a comprehensive tree of life

**DOI:** 10.1101/012260

**Authors:** Cody Hinchliff, Stephen A. Smith, James F. Allman, J. Gordon Burleigh, Ruchi Chaudhary, Lyndon M. Coghill, Keith A. Crandall, Jiabin Deng, Bryan T. Drew, Romina Gazis, Karl Gude, David S. Hibbett, Laura A. Katz, H. Dail Laughinghouse, Emily Jane McTavish, Peter E. Midford, Christopher L. Owen, Richard Ree, Jonathan A. Rees, Douglas E. Soltis, Tiffani Williams, Karen A. Cranston

## Abstract

Reconstructing the phylogenetic relationships that unite all lineages (the tree of life) is a grand challenge. The paucity of homologous character data across disparately related lineages currently renders direct phylogenetic inference untenable. To reconstruct a comprehensive tree of life we therefore synthesized published phylogenies, together with taxonomic classifications for taxa never incorporated into a phylogeny. We present a draft tree containing 2.3 million tips -- the Open Tree of Life. Realization of this tree required the assembly of two additional community resources: 1) a novel comprehensive global reference taxonomy; and 2) a database of published phylogenetic trees mapped to this taxonomy. Our open source framework facilitates community comment and contribution, enabling the tree to be continuously updated when new phylogenetic and taxonomic data become digitally available. While data coverage and phylogenetic conflict across the Open Tree of Life illuminate gaps in both the underlying data available for phylogenetic reconstruction and the publication of trees as digital objects, the tree provides a compelling starting point for community contribution. This comprehensive tree will fuel fundamental research on the nature of biological diversity, ultimately providing up-to-date phylogenies for downstream applications in comparative biology, ecology, conservation biology, climate change, agriculture, and genomics.

## Significance statement

Scientists have used gene sequences and morphological data to construct tens of thousands of evolutionary trees that describe the evolutionary history of animals, plants and microbes. This study is the first to apply an efficient and automated process for assembling published trees into a complete tree of life. This tree, and the underlying data, are available to browse and download from the web, facilitating subsequent analyses that require evolutionary trees. The tree can be easily updated with newly published data. Our analysis of coverage not only reveals gaps in sampling and naming biodiversity, but also further demonstrates that most published phylogenies are not available in digital formats that can be summarized into a tree of life.

## Introduction

The realization that all organisms on Earth are related by common descent (1) was one of the most profound insights in scientific history. The goal of reconstructing the tree of life is one of the most daunting challenges in biology. The scope of the problem is immense: there are ∼1.8 million named species, and most species have yet to be described (2)(3)(4). Despite decades of effort and thousands of phylogenetic studies on diverse clades, we lack a comprehensive tree of life, or even a summary of our current knowledge. One reason for this shortcoming is lack of data. GenBank contains DNA sequences for ∼411,000 species, only 22% of estimated named species. While some gene regions (e.g., *rbcL,* 16S, COI) have been widely sequenced across some lineages, they are insufficient for resolving relationships across the entire tree (5). Most recognized species have never been included in a phylogenetic analysis because no appropriate molecular or morphological data have been collected.

There is extensive publication of new phylogenies, data, and inference methods, but little attention to synthesis. We therefore focus on constructing the first comprehensive tree of life through the integration of published phylogenies with taxonomic information. Phylogenies by systematists with expertise in particular taxa likely represent the best estimates of relationships for individual clades. By focusing on trees instead of raw data, we avoid issues of dataset assembly (6). However, most published phylogenies are available only as journal figures, rather than in electronic formats that can be integrated into databases and synthesis methods (7, 8), (9). Although there are efforts to digitize trees from figures (10), we focus instead on synthesis of published, digitally-available phylogenies.

When source phylogenies are absent or sparsely sampled, taxonomic hierarchies provide structure and completeness (11, 12). Given the limits of data availability, synthesizing phylogeny and taxonomic classification is the only way to construct a tree of life that includes all recognized species. One obstacle has been the absence of a complete, phylogenetically-informed taxonomy that spans traditional taxonomic codes (13). We therefore assembled a comprehensive global reference taxonomy via alignment and merging of multiple openly-available taxonomic resources. The Open Tree Taxonomy (OTT) is open, extensible, and updatable, and reflects the overall phylogeny of life. With the continued updating of phylogenetic information from published studies, this framework is poised to update taxonomy in a phylogenetically-informed manner far more rapidly than has occurred historically.

We used new graph methods (14) to synthesize a tree of life of over 2.3 million OTUs (operational taxonomic units) from the reference taxonomy and curated phylogenies. Taxonomies contribute to the structure only where we do not have phylogenetic trees. Advantages of graph methods include easy storage of topological conflict among underlying source trees in a single database, the construction of alternative synthetic trees, and the ability to continuously update the tree with new phylogenetic and/or taxonomic information. Importantly, our methodology also highlights the current state of knowledge for any given clade and reveals those portions of the tree that most require additional study. Although a massive undertaking in its own right, this draft tree of life represents only a first step. Through feedback, addition of new data, and development of new methods, the broader community can improve this tree.

## Results

### Open Tree Taxonomy

To align phylogenies from different sources, the tips, which may represent different taxonomic levels, must be mapped to a common taxonomic framework (14). For synthesizing phylogenetic data, taxonomy also provides completeness and structure where phylogenetic studies have not sampled all known lineages (true of most clades). Available taxonomies differ in completeness and how closely the hierarchy matches known evolutionary relationships. The Open Tree Taxonomy, OTT, is an automated synthesis of available taxonomies, maximizing the number of taxa and preferring input taxonomies that better align to phylogenetic hypotheses in various clades (see Methods). It contains taxa with traditional Linnaean names and unnamed taxa known only from sequence data. OTT v 2.8 has 2,722,024 OTUs without descendants and includes 382,564 higher taxa; 585,081 of the names are classified as non-phylogenetic units (e.g., *incertae sedis*) and were therefore not included in the synthesis pipeline. The taxonomy is available for download and through web services, including a taxonomic name resolution service for aligning other trees with our taxonomy (see Data and Software Availability, below).

### Input Phylogenies

We built user interface for collection and curation of potential trees for synthesis (https://tree.opentreeoflife.org/curator). The complete database contains 6810 trees from 3062 studies. At the time of publication, 484 studies in our database are incorporated into the draft tree of life. Our goal is to generate a best estimate of phylogenetic knowledge; based on our tests, we give several reasons not to use all available trees for synthesis. First, including trees that are incorrect does not improve the synthetic estimate. In each major clade, expert curators selected and ranked input trees for inclusion based on date of publication, underlying data, and methods of inference (see Methods for details). These rankings generally reflect community consensus about phylogenetic hypotheses. Second, including trees that merely confirm or are subsets of other analyses only increases computational difficulty without significantly improving the synthetic tree. For example, while we have many framework phylogenies spanning angiosperms, we did not include older trees where a newer tree extends the same underlying data. Third, inclusion of trees requires a minimum level of curation, where most OTU labels have been mapped to the taxonomic database, the root is correctly identified, and an ingroup clade has been identified. This information is not in the input file and requires manual curation from the associated publication. Not all trees are sufficiently well-curated; at this point, we have focused curation efforts on trees that will most improve the synthetic tree. The full set of trees in the database is important for other questions such as estimating conflict or studying history of inference in a clade, highlighting the importance of continued deposition and curation of trees into public data repositories. See Dataset 1 for a list of input trees and metadata.

### A draft tree of life

We constructed a tree alignment graph (14), the graph of life, by loading the Open Tree Taxonomy and the 484 rooted phylogenies into a neo4j database. The graph of life contains 2,339,460 leaf nodes (after excluding non-phylogenetic units from OTT), plus 229,801 internal nodes. It preserves conflict among phylogenies and between phylogenies and the taxonomy. To create the synthetic tree, we traverse the graph, resolving conflict based on the rank of inputs, and label accepted branches that trace a synthetic tree summarizing the source information. This allows for clear communication of how conflicts are resolved through ranking, and of the source trees and / or taxonomies that support a particular resolution. The synthetic tree contains phylogenetic structure where we have published trees and taxonomic structure where we do not. See SOM for details. The tree is available to browse and download, and web services allow extraction of subtrees given lists of species (see Data and Software Availability, below).

#### A. Coverage

Of the 2,339,460 tips in the synthetic tree of life, 37,525 are represented in at least one input phylogeny with an additional 4254 non-terminal taxa represented as tips in phylogenetic inputs (Figure 1). In Bacteria, Fungi, Nematoda, and Insecta, there is a large gap between the estimated number of species and what exists in taxonomic and sequence databases (Figure 2). In contrast, Chordata and Embryophyta are nearly fully sampled in databases and in OTT (Figure 2). Poorly sampled clades require more data collection and deposition and, in some cases, formal taxonomic codification and identification to be incorporated in taxonomic databases. Most tips in the synthetic tree are not represented by phylogenetic analyses. The limited number of input trees highlights the need for both new sequencing efforts, additional phylogenetic studies and the deposition of published tree files into data repositories.

**Figure 1:**
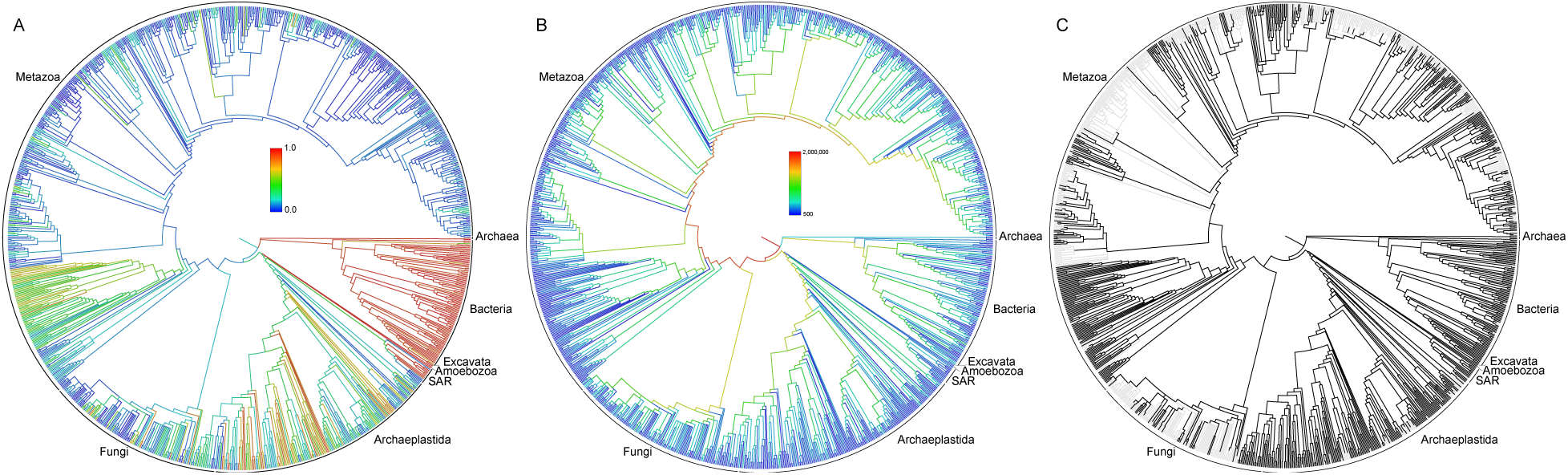
Phylogenies representing the synthetic tree. The depicted tree is limited to lineages containing at least 500 descendants. A. Colors represent proportion of lineages represented in NCBI databases; B. Colors represent the amount of diversity measured by number of descendant tips; C. Dark lineages have at least one representative in an input source tree.

**Figure 2:**
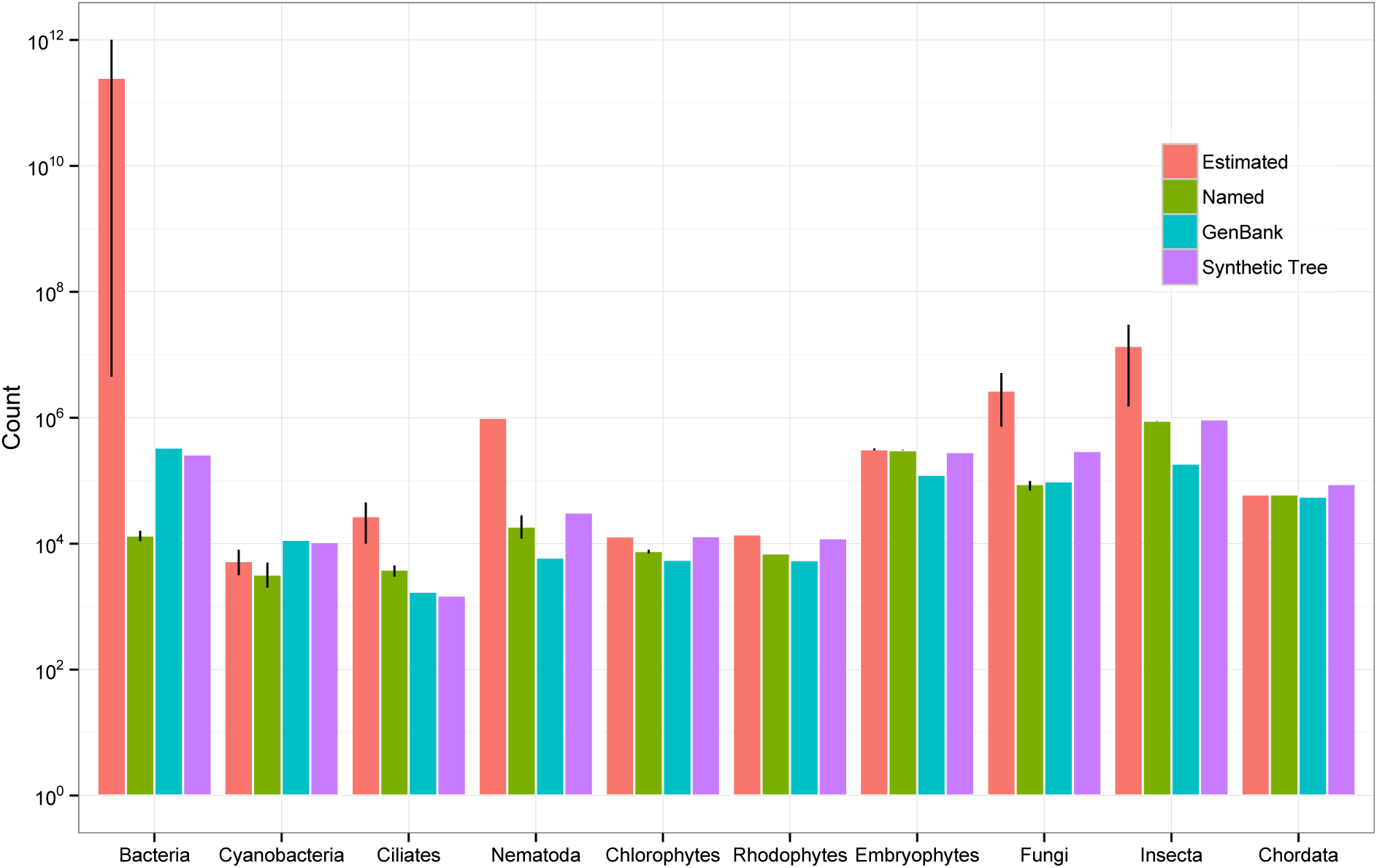
The estimated total number of species, estimated number of named species in taxonomic databases, the number of OTUs with sequence data in GenBank, and the number of OTUs in the synthetic tree, for 10 major clades across the tree of life. Error bars (where present) represent the range of values across multiple sources. See Dataset 2 for underlying data.

#### B. Resolution and conflicts

The tree of life we provide is only one representation of the Open Tree of Life data. Analysis of the full graph database (the graph of life) allows us to examine conflict between the synthetic tree of life, taxonomy, and source phylogenies. Figure 3 depicts the types of alternate resolutions that exist in the graph. We recovered 153,109 clades in the tree of life, of which 129,778 (84.8%) are shared between the tree of life and the Open Tree Taxonomy. There are 23,331 clades that either conflict with the taxonomy (4610 clades; 3.0%) or where the taxonomy is agnostic to the presence of the clade (18721 clades; 12.2%). The average number of children for each node in the taxonomy is 19.4, indicating a poor degree of resolution compared to an average of 2.1 in the input trees. When we combine the taxonomy and phylogenies into the synthetic tree, the resolution improves to an average of 16.0 children per internal node. See SOM for details.

**Figure 3:**
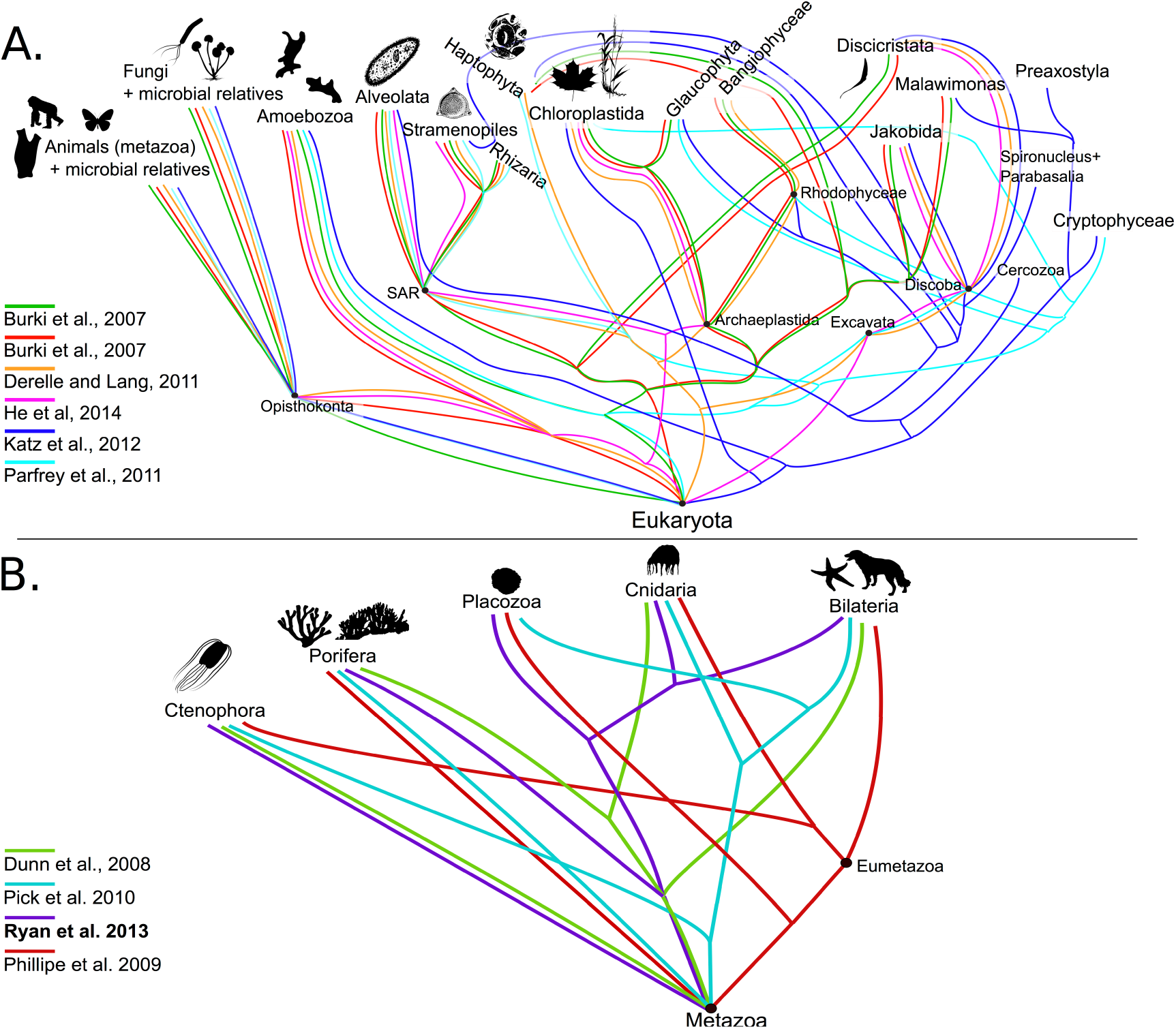
Conflict in the tree of life. While the Open Tree of Life contains only one resolution at any given node, the underlying graph database contains conflict between trees and taxonomy (noting that these figures are conceptual, not a direct visualization of the graph). These two examples highlight ongoing conflict near the base of Eukaryota and Metazoa. Images from PhyloPic (http://phylopic.org).

Alignment of nodes between the synthetic tree and taxonomy reveals how well taxonomy reflects current phylogenetic knowledge. Strong alignment is found in Primates and Mammalia, while our analyses reveal a wide gulf between taxonomy and phylogeny in Fungi, Viridiplantae (green plants), Bacteria, and various microbial eukaryotes (Table 1).

**Table 1:** Alignment between taxonomy and phylogeny in various clades of the tree of life.

| *Clade* | *Tips* | *Internal nodes* | Nodes supported by |  |  |
| --- | --- | --- | --- | --- | --- |
|  |  |  | *Taxonomy* | *Trees* | *Trees + taxonomy* |
| Bacteria | 260323 | 11028 | 8454 (76.7%) | 2184 (19.8%) | 390 (3.5%) |
| Cyanobacteria | 10581 | 788 | 678 (86.0%) | 83 (10.5%) | 27 (3.4%) |
| Ciliates | 1497 | 657 | 654 (99.5%) | 1 (0.2%) | 2 (0.3%) |
| Nematoda | 31287 | 3504 | 3431 (97.9%) | 54 (1.5%) | 19 (0.5%) |
| Chlorophytes | 13100 | 1267 | 1239 (97.8%) | 20 (1.6%) | 8 (0.6%) |
| Rhodophytes | 12214 | 1292 | 1278 (98.9%) | 14 (1.1%) | 0 |
| Fungi | 296667 | 8646 | 8243 (95.3%) | 383 (4.4%) | 20 (0.2%) |
| Insecta | 941753 | 88666 | 85936 (96.9%) | 2205 (2.5%) | 525 (0.6%) |
| Chordata | 88434 | 27315 | 13374 (49.0%) | 11689 (42.8%) | 2250 (8.2%) |
| Primates | 681 | 501 | 129 (25.7%) | 294 (58.7%) | 78 (15.6%) |
| Mammals | 9539 | 4433 | 1645 (37.1%) | 2194 (49.5%) | 594 (13.4%) |
| Embryophytes | 284447 | 32211 | 22400 (69.5%) | 8533 (26.5%) | 1271 (3.9%) |

#### C. Comparison with supertree approaches

There were no supertree methods that scale to phylogenetic reconstruction of the entire tree of life, meaning that our graph synthesis method was the only option for tree-of-life-scale analyses. To compare our method against existing supertree methods, we employed a hybrid MultiLevelSupertree (MLS, (15)) + synthesis approach (see Methods). The total number of internal nodes in the MLS tree is 151458, compared to 155830 in the graph synthesis tree, although the average number of children is the same (16.0 children / node). If we compare the source phylogenies against the MLS supertree and the draft synthetic tree, the synthesis method is better at capturing the signal in the inputs. The average topological error (normalized Robinson-Foulds distance, where 0 = share all clades and 100 = share no clades (16)) of the MLS vs input trees is 31, compared to 15 for the graph synthesis tree. See SOM for details.

## Discussion

Using novel graph database methods, we combine published phylogenetic data and the Open Tree Taxonomy to produce a first draft tree of life with 2.3 million tips -- the Open Tree of Life. This tree is comprehensive in terms of named species, but it is far from complete in terms of biodiversity or phylogenetic knowledge. It does not aim to infer novel phylogenetic relationships, but instead is a summary of published and digitally-available phylogenetic knowledge. This is the first time a comprehensive tree of life has been available for any analyses that requires a phylogeny, even if the species of interest have not been analyzed together in a single, published phylogeny.

As a result of data availability, data quality, and conflict resolution, there are many areas where relationships in the tree do not match current phylogenetic thinking (e.g., relationships within Fabaceae, Compositae, Arthropoda). This draft tree of life represents an initial step. The next step in this community-driven process is for experts to contribute trees and annotate areas of the tree they know best.

### Limitations on coverage

Many microbial eukaryotes, Bacteria, and Archaea are not present in openly available taxonomic databases and therefore not incorporated into the Open Tree Taxonomy and the synthetic tree. Most tips in the synthetic tree (98%) come from taxonomy only, reflecting both the need to incorporate more species into phylogenies and the need to make published phylogenies available. We obtained trees from digital repositories and also by contacting author directly, but our overall success rate was only 16% (8). Many published relationships are not represented in the synthetic tree because this knowledge only exists as journal images. Our infrastructure allows for the synthetic tree to be easily and continuously updated via updated taxonomies and newly published phylogenies. The latter is dependent on authors making tree files available in repositories such as TreeBASE (17), Dryad (http://datadryad.org) or through direct upload to Open Tree of Life (http://tree.opentreeoflife.org/curator) and on having sufficient metadata for trees. We hope this synthetic approach will provide incentive for the community to change the way we view phylogenies - as resources to be cataloged in open repositories rather than simply as static images.

### Conflicts in the tree of life

The synthetic tree of life is a bifurcating phylogeny (with “soft” polytomies reflecting uncertainty), but some relationships are more accurately described using reticulating networks. The Open Tree of Life contains areas with conflict (Figure 3). For example, the monophyly of Archaea is contentious - some data store trees indicate that eukaryotes are embedded within Archaea (18)(19) rather than a separate clade. Similarly, multiple resolutions of early diverging animal lineages have been proposed (20–23). Reticulations help visualize competing hypotheses, gene tree / species tree conflicts, and underlying processes such as HGT, recombination, and hybridization, which have had major impacts throughout the tree of life (e.g., hybridization in diverse clades of green plants (24) and animal lineages (25), including our own (26), and HGT in bacteria and archaea (27–29)). The graphical synthesis approach employed here naturally allows for storage of conflict and non-tree-like structure, enabling downstream visualization, analysis, and annotation of conflict (Figure 3) and highlighting the need for additional work in this area.

Resolving conflict is a challenge in supertree methods, including our graph method. The number of input trees that support a synthetic edge may be considered a reasonable criterion for resolving conflict, but the datasets used to construct each source tree may have overlapping data, making them non-independent. The number of taxa or gene regions involved cannot be used alone without other information to assess the quality of the particular analysis. Better methods for resolving conflict require additional metadata about the underlying data and phylogenetic inference methods.

### Selection of input trees

We used only a subset of trees in the database for synthesis, filtering out trees that are redundant, erroneous or have insufficient metadata. Our current synthesis method relies on manual ranking of input trees by expert curators within major clades. The potential to automate this ranking, and to use metadata to resolve conflict, depends on the availability of machine-readable metadata for trees; such data currently must be entered manually by curators after reading the publication. Additional metadata would allow a comparison of synthesis trees based on, for example, morphological versus molecular data, inference method, or the number of underlying genes. Manual curation is time-consuming and labor-intensive; scalability would improve greatly by having standardized metadata (41) encoded in the files output by inference packages (e.g.,in NeXML files; (30).

### Source trees as a community resource

The availability of well-curated trees allows for many analyses other than synthesis, such as calculating the increase in information content for a clade over time or by a particular project or lab; comparing trees constructed by different approaches; or recording the reduction in conflict in clades over time. These analyses require that tips be mapped to a common taxonomy to compare across trees. Our database contains thousands of trees mapped to existing taxonomies through the Open Tree Taxonomy. The data curation interface is publicly available (http://tree.opentreeoflife.org/curator) as is the underlying data store (http://github.com/opentreeoflife/phylesystem).

### Dark parts of the tree

Hyperdiverse, poorly understood groups including Fungi, microbial eukaryotes, Bacteria, and Archaea are not yet well represented in input taxonomies. Our effort also highlights where major research is needed to achieve a better understanding of existing biodiversity. Metagenomic studies routinely reveal numerous OTUs that cannot be assigned to named species (31, 32). For Archaea and Bacteria, there are additional challenges created by their immense diversity, lack of clarity regarding species concepts, and rampant horizontal gene transfer (HGT) (27)(33)(34). The operational unit is often strains (not species), which are not regulated by any taxonomic code; strain collections are not available to download, making it difficult to map taxa between trees and taxonomy and estimate named biodiversity. Open databases such as BioProject at NCBI (http://www.ncbi.nlm.nih.gov/bioproject) have the potential to better catalog biodiversity that does not fit into traditional taxonomic workflows.

## Materials and methods

### Input data: taxonomy

No single taxonomy is both complete and has a backbone well-informed by phylogenetic studies. We therefore constructed the Open Tree Taxonomy (OTT), by merging Index Fungorum (35), SILVA (36, 37), NCBI (38), GBIF (39), IRMNG (40), and two clade-specific resources (41) (42) using a fully-documented, repeatable process that includes both generalized merge steps and user-defined patches (See SOM). OTT (v 2.8.5) consists of 2,722,024 well-named entities and 1,360,819 synonyms with an additional 585,081 entities having non-biological or taxonomically incomplete names, (“environmental samples” or “incertae sedis”), that are not included in the synthetic phylogeny.

### Input data: phylogenetic trees

We imported and curated phylogenetic trees using a new interface that saves tree data directly into a GitHub repository (43). We obtained published trees from TreeBASE (17) and Dryad, and by direct appeal to authors. The data retrieved are by no means a complete representation of phylogenetic knowledge, as we obtained digital phylogeny files for only 16% of recently published trees (9). Even when available (as newick, NEXUS, NeXML files or via TreeBASE import), trees require significant curation to be usable for synthesis. We mapped taxon labels (which often include lab codes or abbreviations) to taxonomic entities in OTT. We rooted (or re-rooted) trees to match figures from papers. As relationships among outgroup taxa were often problematic, we identified the ingroup / focal clade for the study. For studies with multiple trees, we tagged the tree that best matched the conclusions of the study as “preferred”. Then, within major taxonomic groups (eukaryotic microbial clades, animals, plants and fungi) we ranked preferred trees to generate prioritized lists. In the absence of structured metadata about the phylogenetic methods and data used to infer the input trees, rankings were assembled by authors with expertise in specific clades and were based on date of publication, taxon sampling, the number of genes / characters in the alignment, whether the specific genomic regions are known to be problematic, support values, and phylogenetic reliability (agreement or disagreement with well-established relationships), see Table 2 for details. In general, rankings reflect community consensus about phylogenetic hypotheses. As we collect more metadata - such as that described by the MIAPA, Minimum Information for a Phylogenetic Analysis (44), either by manual entry into the system or by upload of tree files with structured, machine-readable metadata - automated filtering / weighting trees based on metadata will be possible.

**Table 2:** Tree metadata, based on the MIAPA checklist (https://github.com/miapa/miapa). We note whether the metadata is generally available in the tree file (as opposed to in the text of the article, if at all) and how the data is used by Open Tree of Life.

| *Item* | *Description* | *Typically included in tree files* | *Use by Open Tree of Life* |
| --- | --- | --- | --- |
| Topology | The topology itself, plus the type of tree, e.g. gene tree vs species tree, type of consensus tree | Topology, but not tree type | Yes, topology; tree type used by curators as criteria to rank trees |
| Root | Whether the tree is rooted, and the location of the root | Tree in file often rooted arbitrarily; different than in manuscript figures | Yes, requires manual checking by curator to match against manuscript |
| OTU labels | Labels on tips of tree should include (or be mappable to) a meaningful online identifier (e.g. NCBI) | Labels included, but often do not map to online databases due to misspellings, synonymy, etc. | Tip labels mapped through combination of automated and manual processes |
| Branch lengths | The length of each branch of the tree, and the units of measurement | Branch length sometimes included; units generally not present | Imported into database when present but not included on synthetic tree |
| Branch support | Support values (e.g. bootstrap proportions or Bayesian posterior probabilities) | Often in files, but support type often not specified | Not in algorithm, but curators do examine branch support |
| Character matrix | The data used to infer the tree, including data type and source (e.g. GenBank accession or specimen) | Sometimes included with tree file but often without sufficient metadata | Number and type of genes used by curators as criteria to rank trees |
| Alignment method | Method used to align sequence data | No | No |
| Inference method | Method used to infer tree from data | Usually no | Inference method used by curators as criteria to rank trees |

### Synthesis

The goal of the supertree (or “synthesis”) operation is to summarize the ranked input trees and taxonomy (with the taxonomy given the lowest rank). We use an algorithmic approach to produce the synthetic tree rather than a search through tree space for an optimal tree. Given a set of edges labeled with the ranks of supporting trees, the algorithm is a greedy heuristic that tries to maximize the sum of the ranks of the included edges. We summarize the major steps of the method here and provide details in Supplementary Online Materials (SOM).

The first steps include pre-processing the inputs. We prune non-biological or taxonomically incomplete names from OTT, and prune outgroups and unmapped taxa from input trees. Removal of outgroups reduces errors from unexpected relationships among outgroup taxa. Finally, we find uncontested nodes across the taxonomy + input trees and break the inputs at these nodes into a set of subproblems. This allows for a divide-and-conquer approach that shortens running time and reduces memory requirements.

We then build a tree alignment graph (14, 45), which we refer to as the graph of life. Tree alignment graphs allow for representation of both congruence and conflict in the same data structure, allow for non-overlapping taxon sets in the inputs (as well as tips mapped to higher taxa) and are computationally-tractable at the scale of 2.3 million tips and hundreds of input trees. We load the taxonomy nodes and edges into the graph, and then each subproblem, creating new nodes and edges and mapping tree nodes onto compatible taxonomy nodes. We also create new nodes and edges that reflect potential paths between the inputs.

Once the graph is complete, generating the synthetic tree involves traversing the graph and preferring edges that originate from high-ranked inputs. This means that we always prefer phylogenies over taxonomy. Given additional digitized metadata about trees, this system allows for custom synthesis procedures based on preference for inference methods, data types or other factors.

As a comparison to this rank-based analysis, we also created a synthetic tree using MultiLevelSupertrees (MLS) (15), a supertree method where the tips in the source trees can represent different taxonomic hierarchies. We built MLS supertrees for the largest clades that were computationally feasible and then used these non-overlapping trees as input into the graph database and conducted synthesis. Due to the lack of taxon overlap between each MLS tree, there was no topological conflict, and creating the final MLS supertree simply involved traversing the graph and preferring phylogeny over taxonomy.

## Data and software availability

The current version of the tree of life is available for browse, comment and download at https://tree.opentreeoflife.org. All software is open-source and available at https://github.com/opentreeoflife. The tree data store is available at https://github.com/opentreeoflife/phylesystem. Where not limited by pre-existing terms of use, all data are published with a CC-0 copyright waiver. The Open Tree of Life taxonomy, the synthetic tree and processed inputs are available from Dryad: http://dx.doi.org/10.5061/dryad.8j60q

## Acknowledgements

We are grateful to Paul Kirk at Index Fungorum, Tony Rees at IRMNG, and Markus Doering at GBIF for taxonomy data and advice on taxonomy synthesis; Mark Holder for discussion, feedback and software development; Joseph Brown for data collection and curation, software development, data analysis, and writing; Pam Soltis for helpful comments on the manuscript, to authors who made their tree files available in TreeBASE or Dryad and tree files that were not otherwise available, to curators who imported trees and added metadata; and finally to NSF AVATOL #1208809 for funding.

## Supplementary materials and methods

### Introduction

This document contains detail about the construction of the composite taxonomy, curation of input trees, creation of the synthesis tree, and conflict analyses. Fig. S1 gives an overview of the Open Tree of Life process, data stores and services. Constructing the draft tree involves two different types of inputs: taxonomies and published phylogenies. We first combine multiple taxonomic hierarchies into a single taxonomy, the Open Tree Taxonomy (OTT). In a web application built specifically for this project, we input and curate published phylogenies that are then saved to a public GitHub repository.

We load OTT into a neo4j graph database, creating an initial set of nodes and edges. Then, to reduce computational complexity, we decompose the tree inputs into independent subproblems based on nodes that are uncontested across inputs. These subproblems are then loaded into the database, creating a tree alignment graph. Using this data structure, which contains information for all inputs, we extract a tree by resolving conflicts based on a ranked list of inputs, where are trees are ranked higher than taxonomy.

Once we have a draft trees, we assess conflict and compare resolution between the synthetic tree, the taxonomy and the set of input trees.

### Constructing a composite taxonomy

The synthesis of the OpenTree Taxonomy (version 2.8 used here) is a fully automated process. The pipeline takes taxonomy database inputs, in this case, Index Fungorum (17), SILVA (19, 20), NCBI Taxonomy (15), GBIF (16), IRMNG (18) and two clade-specific resources (44, 45). Source taxonomies, each of which is published in its own idiosyncratic representation, are first preprocessed to convert them to a common format. Each source taxonomy in turn is then merged into developing a union taxonomy. Merging a source taxonomy into the union taxonomy consists of two steps: aligning source nodes to union nodes to resolve homonyms, followed by transferring unaligned (new) nodes into the union. A set of about 300 scripted ad hoc manipulations fix errors in the input taxonomies and address situations where automatic alignment has failed. Because the process is scripted, it can be executed any time one of the input taxonomies is revised. The source code for this process is available at https://github.com/OpenTreeOfLife/reference-taxonomy. Version 2.8 of the OpenTree Taxonomy consists of 3,307,105 names, of which 2,722,024 are external (tips) and 585,081 are internal. The taxonomy also has 1,360,819 synonyms. Each name is given a unique id (OTT id) that is used for mapping taxa in trees. The produced taxonomy is then ingested into a neo4j graph database managed by the taxomachine software (https://github.com/OpenTreeOfLife/taxomachine). Taxomachine serves the taxonomy with REST calls over a network and provides a taxonomic name resolution service that allows for disambiguation of taxonomic names as a result of misspellings, changed classification, or homonyms. Taxomachine returns the unique OpenTree taxonomy id for each name in a taxonomic name resolution call.

### Curating input trees

We developed a git-based datastore for phylogenies (46) which connects a web interface (http://tree.opentreeoflife.org/curator) through a programming interface to a GitHub repository (http://github.com/opentreeoflife/phylesystem). The phylogenies in our datastore come from automated import from TreeBASE, input of downloaded files from Dryad, journal supplementary material, and contacting authors directly for files. At time of synthesis, the datastore contained 6753 trees from 3040 studies (see Fig. S2 for distribution of data), although only 484 of these trees are included in this version of the synthetic tree. Studies contained information about the publication, trees, and the list of taxa included. In all cases, the original tree files did not contain sufficient annotation and metadata for synthesis, so there was significant curation by experts. Curation involved two major steps. First, curators mapped the tip labels in the trees to entities in taxonomic databases, assigning an OTT id and disambiguating any problems due to homonyms. Tip labels may be mapped to alternate taxonomic levels (e.g. species, genus, etc.). Second, curators checked that the tree rooting was correct, the ingroup was identified, and that the tree matched the Fig. Sin the publication (tree files deposited in TreeBASE and Dryad often differ from what is shown in the original publication). Ingroups needed to be specified because often the rooting of the outgroup is not accurate, and therefore, relationships in the outgroups may be poor. Once curated, studies are then stored in the phylesystem GitHub repository as Nexson files (NeXML (38) serialized as javascript object notation, JSON). More information about NeXSON can be found at http://purl.org/opentree/nexson.

### Overview of synthesis method

The goal of the supertree (or “synthesis”) operation is to summarize the input trees and taxonomy. We use an algorithmic approach to produce the synthetic tree rather than a search through tree space for an optimal tree. However, one can think of the algorithm as a heuristic attempt to find a tree which maximizes a “number of highly ranked groups displayed” criterion and minimizes unsupported groups. We use a ranking of input trees determined by domain experts to weight groupings.

> *Definition: Number of highly ranked clusters displayed*
>
> By removing parts of the tree, we can restrict a supertree to the *same* set of tips found in any input tree. If a group in the input tree is also found in this pruned form of the supertree, we say that the supertree *displays* the group. If each input tree is assigned a rank, which transfers to all the groupings in that tree, we can summarize the number of input tree groups of each rank that are displayed by a supertree. The supertree that maximizes the number of such displayed groups from highly ranked trees (DCR) would be the preferred tree. Our algorithms attempt to identify a tree (which we call the synthetic tree) that displays as many groups from the top ranked tree as possible, and then displays as many groups as possible from the next most highly ranked tree, *etc*. In order to have an algorithm that can be run in a reasonable time frame, we use a thorough but nonetheless approximate optimization procedure that does not guarantee that we find the best possible tree, but which is computationally efficient and has low rates of error.

> *Definition: “Unsupported” groups*
>
> A distance from the supertree to the inputs can be calculated as a sum of Robinson-Foulds (RF) distances. To calculate an RF distance from a supertree to an input tree, we restrict the supertree to the same set of tip taxa and then calculate the RF distance between this restricted tree and the input tree (49). If a clade in the supertree can be collapsed into a polytomy with its parent group and none of the RF distances from the supertree to the input trees increase, then we say that the clade is *unsupported*. A tree without unsupported groups in this sense is referred to as being a *minimal* tree for the set of groupings that it displays (50).

We use a tree alignment graph (14) to align the input trees and taxonomy and facilitate synthesis. A tree alignment graph (TAG) data structure is a directed acyclic multigraph, containing nodes that represent clade hypotheses and directed edges that represent phylogenetic statements of ancestry and descent. TAGs are designed to contain rooted phylogenetic trees that overlap at compatible nodes. They allow us to store both congruent and conflicting relationships within a single database, along with information about the source data on each edge.

This is an overview of the steps in the synthesis procedure. Details appear in the following sections.

1. Pre-process the inputs trees and taxonomy
2. Decompose the input trees into *uncontested* subproblems
3. Initialize a graph containing nodes and edges from taxonomy
4. Process each subproblem into the taxonomy graph to generate the TAG:
  a. Create nodes and scaffold edges from each subproblem tree fragment
  b. Map tree edges onto corresponding scaffold edges to generate the TAG
5. Identify the set of TAG edges that represent a synthetic tree, using an optimization routine that prefers edges that lead to a higher DCR score.

### Comparison to previous work

We have modified the procedure described in Smith et al (14) in order to remove the dependence on order of the inputs, reduce the number of introduced edges that cannot be directly tied to an input tree or taxonomy, improve computational efficiency, and also improve the ability to accept better resolved clades from lower-ranked trees. Our definition of TAGs differs from that described by Chaudhary et al (48). We have steps to create additional nodes and edges in order to ensure high levels of overlap among compatible input trees, which facilitates more effective synthesis because it enables subtrees from input trees to be grafted onto one another in more possible ways.

### Source tree preprocessing

There are four pre-processing steps that we perform on the input trees and taxonomy before decomposing the inputs into subproblems. We use the treemachine software (https://github.com/OpenTreeOfLife/treemachine) for steps 1 - 3; the otc-prune-taxonomy tool from the C++ library otcetera (https://github.com/OpenTreeOfLife/otcetera) for step 4.

#### Step 1: Prune OTT down to taxonomy for synthesis

Some taxa in the full OTT are flagged as being questionable (“major_rank_conflict”, “major_rank_conflict_direct”, “major_rank_conflict_inherited”, “environmental”, “unclassified_inherited”, “unclassified_direct”, “viral”, “barren”, “not_otu”, “incertae_sedis”, “incertae_sedis_inherited”, “extinct_inherited”, “extinct_direct”, “hidden”, “unclassified”). These questionable taxa are pruned from the taxonomy to produce a taxonomic tree for synthesis. This taxonomic input to synthesis has 2,339,460 terminal taxa, which we refer to as the ‘taxonomy tree’.

#### Step 2: Restrict input phylogenetic estimates to the ingroup

The synthetic tree requires rooted trees, and the position of the root within the outgroup is often uncertain. As part of tree curation, we ask curators to designate an ingroup node, whose descendants are considered part of the tree’s ingroup. When loading the trees, we read in only the ingroup taxa, effectively pruning the outgroup. This reduces errors due to poor taxonomic representation biasing outgroup relationships, and also from the incorrect placement of the root within the outgroup.

#### Step 3: Prune tree tips with missing, ambiguous or nested OTT mappings

Not every curated phylogenetic tree estimate has OTUs that are mapped in manner that is consistent with a clear phylogenetic interpretation. To minimize ambiguity, we first require tips in input trees to be associated with OTT taxa; any tips that are not mapped to OTT taxa are pruned. In cases where *multiple* tips are mapped to the same OTT taxon (for example, population-level sampling), we keep the first encountered tip as an exemplar and prune all others. Finally, when a single tree contains nested mappings—one tip mapped to a genus and another tip mapped to a species in that genus—we use the more nested mapping (e.g. the species rather than the genus) and omit the other mappings.

The post-processed trees are available at http://files.opentreeoflife.org/preprocessed/v3.0/

#### Step 4: Prune the taxonomy to taxa required by phylogenetic inputs

Only a small subset of taxa from the OTT taxonomy are mapped to tips of any of the input phylogenetic trees. 37,325 terminal taxa are mapped to leaves in the input phylogenetic trees and 4,254 non-terminal taxa are mapped to tips of at least one input phylogenetic tree.

If a terminal taxon in OTT occurs only in the taxonomic tree, then the taxon’s final placement in the synthetic tree can be determined using the taxonomy. We prune the taxonomy down to this backbone set of taxa, and used this pruned taxonomy for constructing subproblems that can be solved individually. The pruning step decreases the runtime and memory usage of the subsequent decomposition into subproblems. These subproblems comprise a subset of the leaves found in OTT, but include any terminal taxa for which the phylogenetic inputs provide phylogenetic hypotheses. If a tip of a phylogenetic input is mapped to a non-terminal taxon, then all of the descendants of that taxon are retained when we perform this pruning of the taxonomy. The number of tips in the pruned taxonomy was 127,889, rather than 2,339,460 tips in the portion of OTT used for synthesis.

Note for archiving: pruned taxonomy posted at http://phylo.bio.ku.edu/ot/synth-v3-pruned-taxonomy.tre (see https://github.com/OpenTreeOfLife/otcetera/blob/master/supertree/README.md for instructions on how to regenerate it).

### Decomposition into subproblems

We adopted a divide-and-conquer approach for synthesis to shorten running time and reduce memory requirements. The “divide step” is performed by the otc-uncontested-decompose tool from otcetera. This creates a subproblem for each OTT taxon which is not contested by any input tree. We use the -r command-line flag to otc-uncontested-decompose so that any tip which is mapped to a contested taxon is retained in the input tree (the default behavior is to delete such tips).

> *Definition: Contested taxon*
>
> Consider a taxon (i.e., a clade recognized by taxonomy) *I*_*t*_*|O*_*t*_ (using the notation “ingroup” | “outgroup”) and a fully-resolved phylogenetic tree, *f*. We say that *f* contests the taxon if the most recent common ancestor (mrca) of the tips of *f* which are mapped to the ingroup, *I*_*t*_, is also the ancestor of some tips that are mapped to members of the outgroup, *O*_*t*_. In other words, *f* has at least one grouping that is incompatible with the taxon.
>
> An unresolved tree, *g*, contests the taxon if every fully resolved version of g contests the taxon.
>
> If no input tree contests a non-terminal taxon, we refer to it as an uncontested taxon.

The root node of each subproblem is guaranteed to exist in the final supertree; so the divide step creates constraints on the output. Finding uncontested taxa is easy, and constraining these groups to be monophyletic improves the interpretability of the supertree as a summary of the input trees. The decision to constrain uncontested taxa to be present in the supertree can reduce the total number groups from input trees which are displayed by the supertree. This reduction can happen because some cases of conflict with the taxonomy only arise through the interaction of multiple input trees.

After identifying uncontested taxa, we then generate the set of subproblems. Conceptually this process can be thought of as slicing each of the inputs into fragments of trees, partitioning the fragments into the appropriate subproblem. The tips of the subproblems are mapped either to terminal taxa, to other uncontested taxa, or to a contested taxon that is mapped to one of the tips of a phylogenetic input. Each uncontested taxon is the root of exactly one subproblem. Each uncontested taxon other than the root occurs as a leaf label in exactly one subproblem: the subproblem associated with its least inclusive ancestor which is uncontested.

The details of how the subproblems are created are described in otcetera’s documentation. The following procedure is not the exact algorithm used, but provides a simple explanation that would lead to the same set of subproblems: This procedure of slicing a tree is applied to each input phylogenetic tree (output of preprocessing step #3):

1. Find the the tree’s root taxon - this is the taxon which is the least inclusive common ancestor (LICA) of the taxa associated with the tips of the tree. Map this taxon to the root of the input tree’s root.
2. Perform a tip-to-root traversal over the backbone taxonomy to visit each non-terminal taxa are descendants of the tree’s root taxon.
3. Attempt to map each non-terminal taxon, *X*, to the input tree using the following rules:
  a. If neither *X* nor any of its child taxa are mapped on the tree, then this *X* is not represented in this input tree. Do not map *X* onto this input tree. Move on to the next taxon.
  b. Find the node, *n*, in the phylogenetic tree that is mapped to *X* or is the LICA of all of the taxa contained in *X.* If *n* is also the ancestor of other taxa, then this tree contests the taxon *X*. Do not map *X* onto this tree. Move on to the next taxon.
  c. If *n* is a tip mapped to *X*. Do not map X anywhere else on the tree. Move on to the next taxon.
  d. Otherwise: If the n has out-degree < 2 then will map to the subtending edge of the LICA. If the LICA node has out-degree ≥ 2 and it has not been mapped to a taxon, then *X* will map to this node. Finally, if the LICA node has out-degree ≥ 2 and it has already been mapped to a taxon, then map *X* to the subtending edge of the LICA.
  When we “map *X* to the subtending edge of the LICA” we mean introduce a new node (with out-degree=1) along between the LICA node and its parent.

Fig. S3 shows an example of this process of decorating two input trees with the taxonomic definitions from a taxonomy. In that figure, there are five subproblems, corresponding to the five uncontested (blue) nodes. Each subproblem consists of the nodes from OTT and input trees descended from the uncontested node.

The decomposition procedure then acts by cutting each tree at the uncontested nodes and grouping the resulting fragments by the label of the taxon at the root/breakpoint. The results of the decomposition are shown in Fig. S4. Note that not all input trees need be represented within a subproblem. Many of the phylogenetic statements in the subproblems are trivial (the trees lack any internal, non-root nodes). These trivial statements are retained to make it possible to keep track of which input trees span parts of the synthetic tree.

For the taxonomy and input trees used in this version of the synthetic tree, there are 2792 non-trivial subproblems with between 1 and 15 tree fragments (every subproblem contains a taxonomy fragment).

### Constructing the tree alignment graph

#### Overview of TAG construction

The tree alignment graph (TAG) is a multigraph whose edges are labeled with the source information (input tree or taxonomy). Constructing the TAG takes place in two stages:

1. construct a unigraph with the same nodes as the TAG, which we’ll call the *scaffold,* and
2. extract the TAG from the scaffold by selecting and labeling edges from it.

We load the taxonomy and the subproblems into the scaffold, creating nodes and directed edges representing phylogenetic information. The loading process begins with the taxonomy, creating a scaffold node for every OTT taxon. Then we process each subproblem from the decomposition step. The loading and synthesis procedures are done on each subproblem separately, although all nodes and edges are stored within a single neo4j graph database. In subsequent sections on TAG construction, “input trees” refers to the tree fragments in each subproblem, excluding the taxonomic trees.

During the creation of nodes for each subproblems, we do not create redundant nodes (nodes with identical ingroup and outgroup properties). Much of the complexity of creating the TAG arises from the fact that the input trees only partially overlap with each other in terms of taxonomic content. A single node in an input tree may be represented by a large set of nodes in the TAG, even though the TAG does not contain redundant nodes. However, there may be many nodes in the TAG that have the same ingroup/outgroup properties as an input tree node *when the TAG node’s taxonomic composition is restricted to the taxon set of the input tree*. The loading of input phylogenetic trees includes steps that consider many ways that overlapping trees can overlap and interdigitate. Fig. S5 illustrates the steps involved in TAG creation.

#### Definitions

These definitions are used throughout the TAG creation and synthesis text

##### Ingroup / outgroup property

Each node in the TAG has an “ingroup” property and an “outgroup” property. The ingroup and outgroup are sets of taxa, and the node is intended to be interpreted as a hypothesis that there exists, in nature, a clade that contains (at least) all the taxa in the ingroup and none of the taxa in the outgroup. The intersection of these sets must be empty for every node. The outgroup will be empty in the case of TAG nodes that represent the root of an input tree. We define the functions ingroup(*v*) and outgroup(*v*) for all nodes *v* to be the ingroup (resp. outgroup) of *v*.

##### Merge-compatible nodes

Nodes *p* and *q* are *merge compatible* if no taxa in the outgroup of node *p* are found in the ingroup of node *q*, and vice versa. Any such *p* and *q* make identical phylogenetic statements about the relationships among the taxa they share (but not about taxa they do not share).

##### Nested child of

Consider a pair of nodes *p* and *q,* and the set *w* which is the intersection of the outgroup property of *p* and the ingroup property of *q.* We say that *p* is a nestedchildof *q* if:

1. node *p* and *q* are phylogenetically compatible (there exists a tree *t* that displays both *p* and *q* and with *p* descended from *q*), and
2. *w* is not empty.

This relation is used to generate hierarchies of clade hypotheses that naturally give rise to nested statements of phylogenetic relatedness (similar to a phylogenetic tree or network). The requirement that the outgroup of the child contain at least one element of the ingroup of the enclosing parent ensures that any nested child of some node *q* has the capacity to further resolve relationships among the taxa indicated in the ingroup property of *q*.

An invariant of the TAG and the TAG scaffold is that if there is an edge from *p* to *q*, then *p* is necessarily a nestedchildof *q*.

*Example:* Consider the tree (((*a*,*b*),*c*),*d*). The tree makes various claims about nature (which might not be true), such as the existence of a clade that contains *a* and *b* but not *c* or *d*. When the tree is loaded (see below) we get seven nodes: *abcd*|, *abc*|*d*, *ab*|*cd*, *a*|*bcd*, *b*|*acd*, *c*|*abd*, and *d*|*abc*, where I|O indicates a node with ingroup I and outgroup O. For each edge in the tree, a nestedchildof relationship holds between the nodes corresponding to the head and tail of the edge, for example *ab*|*cd* nestedchildof *abc*|*d*. An edge will be added for each of these nestedchildof relationships.

#### Initializing the scaffold with the taxonomy

We initialize the scaffold data structure by first loading the taxonomy tree from step 1 of the pre-processing, adding a node to the scaffold for each node in the taxonomy. The initial set of edges in the scaffold is the set of edges from the taxonomy (if node *v* in the scaffold represents taxon *t*, an edge will be created to connect *v* to the node which represents the parent taxon of *t*). This set of nodes is added to the scaffold before any of the subproblems are processed.

#### Loading the subproblems

Next we perform the loading procedure on each subproblem independently. Each subproblem contains a set of trees (including a subtree induced from the taxonomy) that span the taxonomic region between the uncontested taxon at the root of the given subproblem and those uncontested taxa at the roots of other subproblems. As each subproblem is loaded, the corresponding taxonomic region is populated with nodes and edges representing the nodes and edges of the subproblem’s input trees. Below we discuss the procedures involved in loading and synthesizing each subproblem itself.

#### Adding nodes for each input tree node

The first step taken when loading a subproblem is to create nodes and edges in the scaffold such that every node and edge in an input tree is represented by a node or edge in the scaffold. We skip the taxonomy subtree when loading each subproblem, as those nodes and edges are already in the scaffold. We also skip trivial trees from an input. For example, if an input tree has only one or two leaves in a subproblem, the group does not add any new nodes. We refer to the set of nodes added in this initial step as *R*.

#### Merging nodes across input trees

Because of the partially overlapping taxonomic sets of the input trees, a node in the optimal supertree might result in multiple different combinations of ingroup/outgroup properties when restricted to the taxonomic set of different input trees. To allow for the possibility that nodes in two different input trees represent the same node in the optimal subtree, we create a set of additional nodes for some of the merge-compatible node pairs in *R*.

We consider adding a “merger node,” *d*, for each pair of merge-compatible nodes, *p* and *q*, where *p* and *q* must originate from different input trees. The ingroup property of *d* is the union of the ingroup properties of *p* and *q*. Similarly, the outgroup of *d* is the union of the outgroup properties of *p* and *q*. Node *d* represents the possibility that the input tree nodes associated with *p* and *q* represent different views of the same node in the synthetic tree of life.

Note that any pair of nodes in *R* which derived from trees with disjunct taxonomic leaf sets could generate a valid merger node. However, this would result in a huge number of new nodes. Hence, we only create new merger nodes when the pair of nodes have at least one ingroup taxon in common.

##### Merger nodes from the root of one tree to nodes in other trees

Let the node representing the root of an input tree fragment *j* be denoted *r*_*j*_. In this step, we consider creating merger nodes between *r*_*j*_ and (non-root) nodes in other trees within the same subproblem. Here, we let *k* denote the index for a different tree (*k* ≠ *j*). We will use *q*_*k*_ to denote the scaffold node that represents *m*_*k*_ where *m*_*k*_ is the shallowest node (the node furthest from the root) in tree *k* for which the three following properties are true:

1. None of the taxa found in tree *j* are found in the outgroup property of *q*_*k*_
2. At least one taxon found in tree *j* is included in the ingroup property of *q*_*k*_
3. Let *ITX* be the least inclusive taxonomy node (from the step 1 version of OTT) that contains all of the tips in tree *j*, and let *OTX* be the least inclusive taxonomy node that contains all of *q*_*k*_’s ingroup and at least one member of *q*_*k*_’s outgroup. Then, we require that *OTX* is not a descendant of *ITX*. This requirement eliminates cycles from the scaffold, whose presence would otherwise interfere with synthesis.

If there is any node *q*_*k*_ satisfying these three conditions, then we create merger nodes for the merger of *r*_*j*_ to the nodes representing each of a set of nodes in tree *k*. Specifically, note that *m*_*k*_ was defined as the shallowest node that satisfies the three properties above, and *q*_*k*_ was the scaffold node that represents it. We will create a merger node from *r*_*j*_ and *q*_*k*_. Additionally for every input tree node, *m*, that is an ancestor of *m*_*k*_, we will create a node for the merger of *r*_*j*_ with the node in the scaffold that represents *m*.

Let *B* be the set nodes added to the scaffold by merging root nodes of one tree to nodes in other trees within the same subproblem.

##### Creation of additional merger nodes

It frequently happens that nodes in two trees have the possibility of mapping to the same node in the synthetic tree. We identify such node pairs and create a third node representing this hypothesis.

For each *ordered* pair of merge-compatible nodes *p* and *q* in two different input tree fragments (which are traversed in preorder) where neither *p* or *q* represent a root node of an input tree fragment, we create a merger node if all of the following conditions are met:

1. *p* and *q* are do not have identical ingroup properties and outgroup properties,
2. *p* and *q* have overlapping ingroup properties,
3. the merger node to be created by merging *p* and *q* proposes a clade that is supported by the phylogenetic information in the input trees (using the definition of node support, below)
4. no ancestor of *p* in its input tree has already been used to create a merger node with any node in the input tree containing *q*, and no ancestor of *q* in its input tree has already been used to create a merger node with *p*—this pair of conditions ensures that we only create opportunities for trees to overlap at the most conservative (i.e. deepest) locations within them.

Let *D* be the set of nodes added to the scaffold by merging non-root nodes in input trees.

##### Node support

Creating all possible merger nodes leads to possibility of introducing edges in the synthesis tree that are not supported by input trees. For each potential merger node, we therefore assess whether there is sufficient information in the input trees to separate all the taxa in the proposed merger node’s ingroup from all the taxa in its outgroup, and only create TAG nodes for merger nodes that pass the test. The goal of this ‘sum test’ is to eliminate nodes from the scaffold that could produce edges in the synthetic tree that do not pass the minimal tree test *sensu* Semple (50). The sum test itself is actually more stringent than Semple’s minimal tree test, and may also eliminate some potential edges from the synthesis tree that the latter would not find to be unsupported. However, the minimal tree test is designed to be used over the edges of a single tree, and cannot be used directly to test individual merger nodes before they are added to the TAG, while the sum test can.

The sum test determines if, given a potential merger node *x*, there exists some set *R* of rooted triples where each triple in *R* is displayed by some input tree, such that for every rooted triple *t* implied by *x*, *t* is either contained in or induced by *R*, according to the second definition of ‘induces’ in Guillemot and Berry (51). The goal of this test is to assess whether or not there exists sufficient phylogenetic information in the source trees to imply the separation of all the taxa in *x*’s ingroup from all the taxa in *x*’s outgroup.

##### Edges resulting from the addition of merger nodes

When creating a merger node *d* from nodes p and q, if p or q has has an edge to or from another node x (a parent or child), then a scaffold edge is added between d and x as long as the nestedchildof relation still holds between d and x. These edges allow paths to cross from one input tree to another in the scaffold.

##### Edges linking input trees

As a side effect of the search for merger nodes, additional nestedchildof relationships are identified that link input trees. For each pair of nodes *p* and *q* from different input trees, an edge from *p* to *q* is added to the scaffold whenever *p* is a nestedchildof *q*. This exhaustive set of edges contrasts with the situation where the input trees are added to the scaffold (above).

#### Adding nodes by walking paths in the scaffold

In the merger nodes step, we create only a small subset of the possible set of nodes that can be created by combining nodes from input trees. In the WalkPaths procedure, we consider another way of combining nodes and use it to add more nodes to the scaffold. Let *E* be the set of nodes created for this subproblem thus far. *E* is the union of *R* (the nodes coming directly from input trees), *B* (the set of nodes added by merging root nodes of input trees), and *D* (the set of nodes added by merging non-root nodes).

The nodes created in the previous subproblem-specific sections (the members of the sets *R*, *B* and *D*) have ingroup and outgroup properties that each reflect either 1 or 2 input trees. Members of *R* represent nodes in single trees. Members of *B* were created by merger of root nodes from one tree with a node of a different tree. So, *E* is not exhaustive in the sense that the ingroup+outgroup properties of all of the nodes in the optimal summary for this subproblem may not be represented.

To consider a wider range of possible internal nodes in the subproblem solution, the next step creates nodes with the ingroup+outgroup properties that would be obtained by walking a path connecting elements in *E*. The WalkPaths procedure considers possible paths through these nodes, and produces “accumulation nodes” for the path. Each accumulation node is produced by a node, *x*, along the path. The ingroup property of the accumulation node will be the union of the ingroup properties of *x* and all of the previous nodes in the path. The outgroup property of the accumulation node will be the union of the outgroup properties of *x* and all of the subsequent nodes in the path.

A scaffold node can be proposed if the next node in a path is compatible with the accumulated ingroup property that has been built while traversing the path up to that point.

The WalkPaths procedure starts at each node in *E* and tries to extend that path to a new node consistent with the requirements that:

1. each node in the path is the nested child of the next node, and
2. each node’s outgroup has no overlap with the cumulative ingroup of the previous nodes in the path.

An accumulation node is created for each node in a valid path that has been extended to more than one node.

##### Edges resulting from the addition of accumulation nodes

Three kinds of edges are added to the scaffold with the introduction of each accumulation node *c*.

1. Since *c* derives from path node *x*, we duplicate the edges going in and out of *x*, just as we did for merger nodes.
2. Edges are created between *c* and every node in the path containing *x* (other than *x*).
3. Edges are also created between *c* and every other upstream and downstream accumulation node in its path.

#### Deriving the TAG from the scaffold

With the completion of the scaffold it is possible to say what the TAG is. The nodes of the TAG are simply the nodes of the scaffold, and every edge of the TAG represents a nestedchildof relationship between two nodes that is present as an edge in the scaffold. For synthesis purposes it is important to know which input tree is associated with each edge in the TAG, so every edge in the TAG is labeled with the associated tree and the corresponding edge in the input tree. Scaffold edges that cannot be associated with input trees are not added to the TAG. Because there can be many trees associated with a single scaffold edge, the TAG is a multigraph. The TAG will have an edge (*v*, *w*) labeled with tree *t* and edge (*x, y*) if and only if *t* has an edge (*x, y*) corresponding to (*v*, *w*), where “corresponding to” is determined by the procedure described below.

It might seem better to prevent the creation of non-corresponding edges in the scaffold in the first place, but this check is not easy to perform, so it is left for batch processing at this point.

To determine the edges of the TAG, the edges of each input tree *t* are traversed in preorder, and their corresponding scaffold edges are found. To help find the scaffold edges corresponding to input tree edges, we keep track of a set of scaffold nodes corresponding to each input tree node *x*; call this set *m*(x). The traversal is initialized with all higher taxon nodes within the subproblem which contain all the terminal taxa identified as tips in the tree, i.e. *m*(root(*t*)) = all taxonomy nodes *u* such that ingroup(root(*t*)) ⊆ ingroup(*u*).

Let (*x, y*) be the next edge in the input tree visited in the traversal, with *y* = parent(*x*). Then the edges (*v*, *w*) in the scaffold corresponding to (*x, y*) are those for which the following hold:

- *w* ∈ *m*(*y*)
- ingroup(*x*) ⊆ ingroup(*v*)
- outgroup(x) ⊆ outgroup(v)
- *v* nestedchildof *y*

Each edge (*v*, *w*) from the scaffold corresponding to some edge (*x, y*) in an input tree *t* gives rise in the TAG to an edge (*v*, *w*) with labels *t* and *d*. The traversal continues with *m*(*x*) equal to all nodes *v* for which (*x, y*) has a corresponding scaffold edge (*v*, *w*) as above.

Having now produced a TAG, the scaffold is no longer needed.

### Generating the synthetic tree

Once the TAG has been produced, we use an optimization procedure to find a tree, consisting of edges in the TAG, that generally prefers edges from high ranked trees to edges from lower ranked trees. The optimization works locally: For each TAG node, it selects one “best” combination of edges, from among all possible combinations, that would become the children of the node were that node to be included in the final synthetic tree. See Fig. S6 for an overview.

For each TAG node *v*, let *E*(*v*) be the set of TAG edges that have *v* as their parent node. For each *v* we select a subset *R*(*v*) of *E*(*v*) having the following two properties:

1. *R*(*v*) is a *candidate* subset, i.e. if *v*_*1*_ and *v*_*2*_ are child nodes of edges in *R*(*v*), then no taxon reachable from *v*_*1*_ is reachable from *v*_*2*_. That is, the children of edges in *R*(*v*) are the root nodes of a non-overlapping forest of synthetic subtrees.
2. *R*(*v*) is a “best” candidate subset of *E*(*v*) under the preference ordering, defined below.

The preference ordering on edge sets (which is consistent with the DCR score discussed above) is as follows: Given a candidate edge set *S*(*v*) and an alternate candidate edge set *S*alt(*v*), we say that *S*(*v*) is *better than S*alt(*v*) if, for some rank *d*, it contains more edges with rank *d* and contains an equal number of edges of any rank *f* > *d* where *f* is a rank represented by any other edge in *E*(*v*). (An edge has rank *d* if the tree it is labeled with has rank *d*). If two candidates contain the same number of edges for all ranks in *E*(*v*), then we say that the candidate defining the forest of subtrees that contains the most tip nodes is better, and if they contain the same number of edges of each rank and the same number of tips then we say that they are *equal*. A *best candidate S*max(*v*) is any candidate which is better than or equal to all other possible candidates *S*(*v*). If there is more than one best candidate for some node, then one is selected arbitrarily.

The synthetic tree is the tree *Tr*(*r*) where the root of the tree *r* is the TAG node specified to become the root. That is, *r* is supplied as an input to the synthesis procedure.

Due to conflicts among input trees, it is possible for some tips (taxa) to be left out of the resulting synthesis tree. Those tips are added back into the tree using the procedure described in the next section.

#### Adding missing terminal taxa

The final stage of synthesis involves the attachment of terminal taxa that could not be included by the main synthesis procedure discussed above. When a decision is made during synthesis to prefer a path that excludes a given node, then that contested node will not be included in the synthetic tree. When all the paths to some terminal taxon pass through such contested nodes, then that terminal taxon will likewise not be included in the synthetic tree as it is generated by the procedure described in the previous section. We call such missing terminal taxa ‘missing children’.

Attaching these missing children to the synthetic tree thus requires the creation of additional TAG edges. Attachment of a missing child *y* occurs at the node in the synthetic tree closest to the tips that contains in its ingroup all taxonomic sister taxa of *y*. For example, if *Pseudacris crucifer* were not included in the tree, it would be attached to the most derived TAG node already present in the synthesis tree which serves as an ancestor for all *Pseudacris* species (*P. feriarum*, *P. regilla,* etc.). More formally, given y and immediate taxonomic parent *z*, find the full set *u*_*i*_ of taxonomic terminal child nodes of *z* as well as the set *x*_*i*_ of taxonomic terminal child nodes of *z* that are already present in the synthetic tree (which will be a subset of *u*_*i*_). The TAG node *w* which represents the mrca of taxa *x*_*i*_ in the synthesis tree serves as the first potential attachment point of *y*. If node *w* contains in its ingroup all of *u*_*i*_, it serves as an appropriate attachment point of *y*. However, this need not be the case: for example, if terminal taxa *u*_*i*_ differ in their taxonomic nestedness below *z* (say, some taxa belong to a subgenus while others do not), the mrca node *w* may encompass a restricted set of *u*_*i*_ (i.e. not including *y*). In this case, subsequent rootward nodes in the synthesis tree are tested until the first node that passes this condition is encountered. Performing this procedure ensures that all terminal taxa are present in the final synthetic tree.

### Caveats to the synthesis method

The current methods for generating the first draft synthetic tree were designed to be computationally tractable at the very large scale that the tree of life represents, and their aim is to provide a result (the synthetic tree) that is a good approximation of the best possible tree summarizing the inputs. Here we list known issues and caveats. For all of these problems, we are either currently pursuing, or anticipate future work that will provide solutions.

- The process by which we create nodes and edges in the graph makes concessions to reduce computational complexity (see Merger Nodes and Walking Paths sections above), and as a result may not find all possible paths (i.e. possible edges in the synthesis tree) among nodes in the input trees and taxonomy.
- Polytomies in input trees where not all children of the polytomy are tips are currently handled as “hard polytomies”, and will be maintained in the synthetic tree even if a lower ranked tree has the ability to resolve them.
- When a tip node in a tree is mapped to higher taxon and no higher-ranked tree conflicts with the monophyly of that taxon, that taxon will be displayed as monophyletic in the synthetic tree, even if a lower ranked tree contests the monophyly of that taxon.
- Because we do not perform exhaustive support checks for nodes added to the scaffold during the Merger Nodes and Walk Paths steps, it is possible for the TAG (and the synthetic tree generated from it) to contain nodes that may not be supported according to some provided definition of support. The otcetera program implements Semple’s minimal tree support criterion (50), and finds 9 nodes in the current version of the synthetic tree for which this test finds no support. There are 229,801 total internal nodes in the tree, providing a total unsupported node rate of < 0.00004.
- The procedure to add taxa back to the synthesis tree that have been excluded due to conflict among input trees (see Adding missing terminal taxa section) can rarely place taxa in the synthetic tree in locations that do not correspond with taxonomy. This results in nodes in the synthetic tree that are labeled with a taxonomic name but which do not contain all the descendants of the original taxon in OTT. There are 2 such nodes in the current version of the synthetic tree out of 203,466 non-terminal taxon nodes, yielding an taxonomic mismatch rate of < 0.00001. We are working to resolve this problem.

### Assessing conflict

We measured support for the nodes in the supertree using an approach described by Wilkinson et al. (52). Let c be a clade in the supertree S and c’ be its restriction to the leaves of an input tree T, i.e., c’ contains only those leaves that are present in T. If c’ contains all the leaves of T or less than 2 leaves, then T is *irrelevant* to c. T *supports* c if c’ is present in T. T *conflicts* with c when the induced bipartition (of c) contradicts the relationships in T (16). T *permits* c if c’ is a resolution of a polytomy in T, thus T is agnostic with respect to c. See Fig. S7 for an example.

First, we compared the taxonomy tree to the Open Tree of Life. There are 155,830 clades in the Open Tree of Life, and 129,502 (83.1%) of these are supported by the taxonomy tree. There are 5,340 (3.4%) clades in the Open Tree of Life that are in conflict with the taxonomy, and 20,988 (13.47%) that are permitted.

When we compare the collection of 484 non-taxonomy input trees to the Open Tree of Life, there are 30,550 (19.6%) clades in the Open Tree of Life that are unambiguously supported (i.e., ≥1 non-taxonomy input trees support and 0 non-taxonomy input trees conflict with or permit the node) and 589 (0.4%) clades are in unambiguous conflict (i.e., ≥1 non-taxonomy input trees conflict with and 0 non-taxonomy input trees support or permit the node). However, 123,346 (79.2%) of the clades in the Open Tree of Life are irrelevant to the non-taxonomy input trees. Thus, the information for most of the clades in the Open Tree of Life is coming from the taxonomy. The remaining 1345 (0.9%) nodes in the Open Tree of Life have a combination of support, conflict, and permit, instead of complete support or conflict, among the non-taxonomy input trees with respect to these clades. Overall, 3286 clades in the Open Tree of Life are supported by at least two non-taxonomy input trees. When we compile all of the input trees together (the taxonomy tree and the 484 non-taxonomy input trees), there are 149,567 (96.0%) nodes in the Open Tree of Life that are unambiguously supported by all relevant input trees and 578 (0.4%) clades are in unambiguous conflict. None of the all assessed clades was irrelevant to the input trees. The remaining 5685 (3.7%) nodes in the Open Tree of Life have a combination of support, conflict, and permit, instead of complete support or conflict, among the non-taxonomy input trees with respect to these clades. Overall, 8005 clades in the Open Tree of Life are supported by at least two taxonomy or non-taxonomy input trees.

In contrast, when we compare the collection of 484 non-taxonomy input trees to the MLS Tree of Life, there are 24,891 (16.4%) clades in the MLS Tree of Life that are unambiguously supported (i.e., ≥1 non-taxonomy input trees support and 0 non-taxonomy input trees conflict with or permit the node) and 2,013 (1.3%) clades are in unambiguous conflict (i.e., ≥1 non-taxonomy input trees conflict with and 0 non-taxonomy input trees support or permit the node). However, 123,330 (81.4%) of the clades in the MLS Tree of Life are irrelevant to the non-taxonomy input trees. The remaining 1,224 (0.8%) nodes in the MLS Tree of Life have a combination of support, conflict, and permit, instead of complete support or conflict, among the non-taxonomy input trees with respect to these clades. Overall, 2,693 (1.8%) clades in the MLS Tree of Life are supported by at least two non-taxonomy input trees.

The software for conflict analysis is available at https://github.com/ruchiherself/AssessSupertrees.

### Supplementary Fig. Captions

**Fig. S1: The Open Tree of Life workflow.** External taxonomies (and synonym lists) are merged into the Open Tree Taxonomy, OTT. Published phylogenies are curated (rooted, and names mapped to OTT) and stored, with full edit history, in a GitHub repository. The source trees are decomposed into subproblems, and the loaded along with OTT into a common graph database. We traverse the resulting graph and extract a tree of life based on priority of inputs. Components with stars in indicate presence of application programming interfaces (APIs) to access data and services.

**Fig. S2: Size and scope of input trees.** Plot of the number of tips in each of the 1188 trees with some curation in the treestore. Scope is measured as the total number of tips recognized to be descended from the inferred most recent common ancestor of the source tree.

**Fig. S3: Inputs to subproblem decomposition with taxonomic mappings added**. Uncontested taxa are shown in blue, and contested taxa are shown in red. The hollow circles at nodes in the phylogenetic trees represent internal nodes in the tree that do not map to any taxon. Note that uncontested taxon will not map to the taxon that contests it. This example generates five subproblems, one for each uncontested node.

**Fig. S4: Decomposition into subproblems.** The output of the decomposition into subproblems from the inputs shown in Fig. S3. Some nodes with outdegree=1 have been suppressed, as they are not needed in the rest of the pipeline. The node colorations in this Fig. Sare only retained to make it easier to compare the outputs to the inputs in the previous figure; the status of a node as contested does not matter for the rest of the pipeline.

**Fig. S5: Creation of the tree alignment graph (TAG)**. We initialize the graph with nodes and edges for the taxonomy. Then, we create the graph nodes during the ‘merger nodes’ and ‘accumulation nodes’ steps, and add scaffold edges which identify a subset of nested child relationships among the nodes. Finally, we map the input tree edges onto corresponding edges in the scaffold, creating the TAG (colored edges in the final graph in lower right).

**Fig. S6: Generating the synthetic tree from the TAG.** In synthesis, the nodes of the TAG are visited in topological order. At each node, a decision is made about which child edges would be included as child branches of the node, if the node were to occur in the final synthesis tree. Because nodes are visited in topological order, when we visit some node *x*, decisions have already been made for each child of *x* (and each of their children, and so forth), which means that each child node of *x* is the root of a synthesis subtree defined by those edges that have been selected at all of its descendant nodes. In other words, the procedure to select child edges to include at a given node can be thought of as a procedure to select the subtrees that would be the children of *x* in the final synthesis tree. The decision regarding which edges (i.e. subtrees) to include uses the DCR criterion—that is, the selected subtrees are those which contain the most TAG edges corresponding to edges in highly weighted input trees. However, no two subtrees may be included which contain any tips in common, as this would define a network rather than a tree. In this example, TAG edge colors identify source trees (corresponding to Fig. S5), and colored numbers identify the corresponding source tree edges. Each source tree edge corresponds to at least one edge in the TAG. At each node, a decision is made which edges would be included, which defines a set of unique input tree edges that would be represented in the synthesis subtree below the given node (shown in the list on the lower left). Any edges that are parallel to selected edges are also considered to be represented. The synthesis decision at the root (node 1) selects the edges leading to nodes 7 and 11, and rejects the edges leading to nodes 9 and 11, because this maximizes the representation of tree edges in the final synthesis tree. Note that input tree edges leading to tips (i.e. external edges) are considered represented if the tip itself occurs in the synthesis tree, regardless of whether the specific edge in question does or not.

**Fig. S7: Conflict analysis.** A supertree S with two internal nodes u and v, and three input trees T_1_, T_2_, and T_3_. The clade u is in conflict with T_1_, supported by T_2_, irrelevant to T_3_. The clade v is irrelevant to T_1_ and T_2_, and permitted by T_3_, as v is a resolution of the polytomy at the root in T_3_.

